# Smooth Muscle Cells-Expressed Bmal1 Regulates Vascular Calcification Independent of the Canonical Circadian Pathway

**DOI:** 10.1101/2024.07.22.604443

**Authors:** Ming He, Yong Sun, Fengyuan Huang, Zhehao Zhu, Erandi Velazquez-Miranda, Zechen Chong, Yabing Chen

## Abstract

Vascular calcification is a major cardiovascular issue that increases morbidity and mortality in diabetes patients. While dysregulation of the circadian master regulator Basic Helix-Loop-Helix ARNT-Like Protein 1 (Bmal1) in vascular smooth muscle cells (VSMC) under diabetic conditions has been suggested, its role in vascular calcification is unclear. In VSMC, Bmal1 was upregulated under high glucose treatment and in aortic tissues from a diabetic mouse model. RNA sequencing from isolated VSMC between Bmal1 deletion and wildtype mice indicated Bmal1’s pro-calcification role. Indeed, reduced levels of the osteogenic master regulator, Runt-Related Transcription Factor 2 (Runx2), were found in Bmal1 deletion VSMC under diabetic conditions. Alizarin red staining showed reduced calcification in Bmal1 deletion VSMC *in vitro* and vascular rings *ex vivo*. Furthermore, in a diabetic mouse model, SMC-Bmal1 deletion showed reduced calcium deposition in aortas. Collectively, diabetes-upregulated circadian regulator Bmal1 in VSMC contributes to vascular calcification. Maintaining normal circadian regulation may improve vascular health in diabetes.

---

Diabetes-derived cardiovascular complications are a major cause of mortality, accounting for nearly half of diabetes-related deaths^1^. Hyperglycemia leads to disruptions in cardiovascular function, resulting in impaired cardiac contractility, vascular cell dysfunction, and vascular calcification^2^. Vascular calcification is characterized by the trans-differentiation of vascular smooth muscle cells (VSMC) from a contractile phenotype into an osteoblast-like phenotype after exposure to various stimuli, such as oxidative stress and high glucose levels^3^. Circadian rhythms are known to regulate glucose metabolism under physiological conditions; however, diabetes disturbs circadian rhythms, exacerbating hormonal imbalances^4^. Although there is increasing evidence linking hyperglycemia and circadian disruption in diabetes and cardiovascular diseases^5^, the molecular mechanisms and pathological consequences remain incompletely understood. Here, we identified the dysregulation of the circadian master regulator Basic helix-loop-helix ARNT-like protein 1 (Bmal1) in VSMC under a high-glucose environment. Bmal1 upregulates Runt-related transcription factor 2 (Runx2) and induces VSMC to transition toward an osteogenic phenotype, ultimately leading to increased vascular calcification. Importantly, this regulation in VSMC operates via novel mechanisms independently of traditional circadian regulation pathways. To explore the involvement of circadian proteins in VSMC under diabetic conditions, isolated VSMC from wildtype mice were treated with a medium containing 4.5 g/L glucose. As a control, a medium containing 1 g/L glucose was used. As shown in Panel A, circadian master regulator Bmal1, but not Clock or Bmal2, was induced by high glucose in VSMC. Similarly, Bmal1 was significantly upregulated in aortic tissues from a streptozotocin (STZ)-induced diabetic mouse model, as assessed by Western blot analysis (Panel B). To elucidate the biological function of Bmal1 in VSMC, we generated a smooth muscle cell (SMC)-specific Bmal1 deletion mouse model by breeding the Bmal1^flox/flox^ (Bmal1^f/f^) mice with SM22α-Cre transgenic mice (Panel C).

RNA sequencing analysis revealed significant alterations in the transcriptomes between Bmal1^f/f^ (wildtype) and Bmal1^Δ/Δ^ (knockout) VSMC. Among the 1472 differentially expressed genes (with a fold change greater than 1.5 and a false discovery rate [FDR] less than 0.1), 496 genes were downregulated in the Bmal1^Δ/Δ^ VSMC (Panel D). Gene Ontology analysis further identified Bma1-depeendent signaling pathways in VSMC. In additional to its known function in “E-box binding”, Bmal1-induced DEGs were also involved in “regulation of smooth muscle cell proliferation”, “inflammatory response”, “bone trabecula formation”, “extracellular matrix organization”, “regulation of osteoblast differentiation”, “collagen fibril organization”, “cardiac epithelial to mesenchymal transition”, “cellular response to TGFβ stimulus”, and “cellular response to reactive oxygen species”, suggesting a potential pro-osteogenic role for Bmal1 in VSMC (Panel E). Consistent with RNA sequencing analysis, the expression of the master osteogenic transcription factor, Runx2, and its downstream genes [Osteocalcin (OC)], both induced by diabetic conditions, were diminished at the protein levels in Bmal1 deficient VSMC (Panel F). Functionally, Alizarin Red staining demonstrated significantly reduced calcification in Bmal1 deficient VSMC (Panel G). *Ex vivo* experiments using aortic arteries isolated from SMC-specific Bmal1 deletion mice showed markedly reduced vascular calcification-induced by diabetic conditions, when compared with those arteries isolated from their control littermates (Panel H). Using the low-dose STZ-induced diabetic mouse model, we observed that SMC-specific Bmal1 deletion did not alter blood glucose levels or body weight, while attenuating STZ-induced vascular calcification, as evidenced by total calcium measurement in the descending aortas. This phenotype was consistent in both male and female mice (Panels I).

Collectively, our studies have demonstrated that hyperglycemia associated with diabetes promotes the expression of the circadian regulator Bmal1, which mediates diabetes-induced vascular calcification *via* a novel mechanism through the regulation of Runx2 in VSMC, independently of other circadian proteins. This non-canonical circadian pathway may be specific to the pathogenesis of vascular calcification in diabetes. Of note, our studies also showed that disruption of circadian proteins (e.g., Bmal1 deletion in VSMC) does not systematically alter glucose metabolism. However, excessive Bmal1 in VSMC negatively impacts vascular health. Therefore, maintaining a normal circadian regulation or adjusting dysfunctional circadian factors, like Bmal1, could be a feasible approach to improving vascular health in diabetes and other risk conditions for cardiovascular diseases.

## Acknowledgements

We thank Dilani Patel and Jing Jiang at UAB for technical assistance.

## Figure and Figure Legend

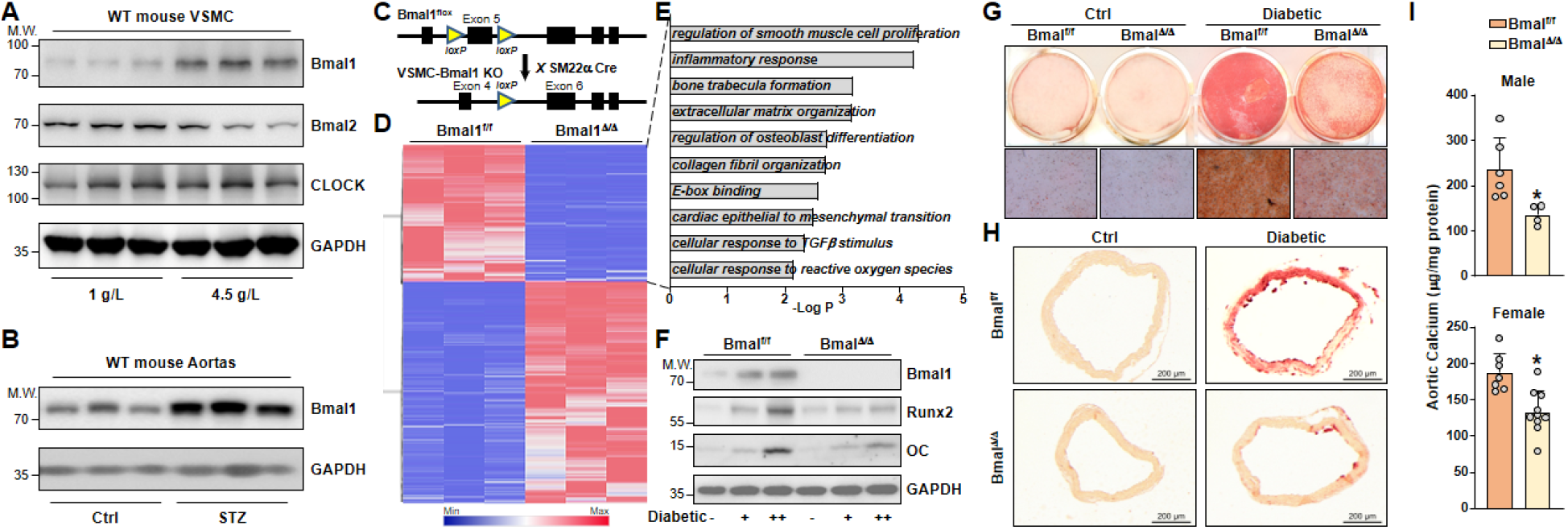
VSMC Bmal1 regulates diabetes-induced vascular calcification. (**A**) Isolated wild-type (WT) mouse vascular smooth muscle cells (VSMC) were cultured in growth medium containing either 1 g/L or 4.5 g/L glucose for 72 hours. (**B**) WT mice were injected a low dose streptozotocin (STZ, 50 mg/kg body weight/day) or control solvent intraperitoneally for 5 consecutive days. Descending aortas were isolated for Western blot analysis 4 months later. (**C**) A schematic diagram illustrates the generation of smooth muscle cell (SMC)-specific Bmal1 deletion mice. (**D**) Heatmap comparison of log2 fold change (FC) for differentially expressed genes in VSMC isolated from Bmal1^f/f^ and SMC-specific Bmal1 deletion (KO, or Bmal1^Δ/Δ^) mice (FC > 1.5, FDR < 0.1). (**E**) Gene Ontology enrichment analysis using Metascape identified top down-regulated signaling pathways in the Bmal1 KO VSMC. (**F**-**G**) VSMC isolated from Bmal1^f/f^ and SMC Bmal1 KO mice (KO, or Bmal1^Δ/Δ^) were cultured under diabetic conditions [osteogenic medium containing 4.5 g/L glucose with 0.15 mM (+) or 0.3 mM (++) H_2_O_2_] or control conditions (growth medium, labeled – or Ctrl). In (F), Bmal1, Runx2, osteocalcin (OC), and GAPDH were detected by Western blot. In (G), VSMC calcification was evaluated using Alizarin Red staining. (**H**) Descending aortic arteries were isolated from mice and cultured in diabetic conditions or control conditions. *Ex vivo* vascular calcification was detected by Alizarin Red staining. (**I**) Male and female Bmal1^f/f^ and SMC-specific Bmal1 deletion mice were injected with low-dose STZ injections as described above. After 4 months, descending aortic arteries were isolated and total calcium content was assessed by the Arsenazo method. The University of Alabama at Birmingham (UAB) animal care personnel maintained all animals in accordance with NIH guidelines, and the UAB Institutional Animal Care and Use Committee (IACUC) approved all experimental procedures (approval No. IACUC-21647). In (I), data are expressed as mean ± SEM. Data were initially tested for normality and equal variance to confirm the appropriateness of parametric tests. If the normality or equal variance test failed, nonparametric data were analyzed using a 2-tailed Mann-Whitney *U* test. A significance level of *p* < 0.05 was considered statistically significant.

## References

1. Sridharan Raghavan, Jason L Vassy, Yuk-Lam Ho, Rebecca J Song, David R Gagnon, Kelly Cho, Peter W F Wilson, Lawrence S Phillips. Diabetes Mellitus-Related All-Cause and Cardiovascular Mortality in a National Cohort of Adults. J Am Heart Assoc. 2019;8(4):e011295.

2. Christian Rask-Madsen, George L King. Vascular complications of diabetes: mechanisms of injury and protective factors. Cell Metab. 2013;17(1):20–33.

3. Yabing Chen, Xinyang Zhao, Hui Wu. Arterial Stiffness: A Focus on Vascular Calcification and Its Link to Bone Mineralization. Arterioscler Thromb Vasc Biol. 2020;40(5):1078–1093.

4. Zhilei Shan, Hongfei Ma, Manling Xie, Peipei Yan, Yanjun Guo, Wei Bao, Ying Rong, Chandra L Jackson, Frank B Hu, Liegang Liu. Sleep duration and risk of type 2 diabetes: a meta-analysis of prospective studies. Diabetes Care. 2015;38(3):529–37.

5. Ivy C Mason, Jingyi Qian, Gail K Adler, Frank A J L Scheer. Impact of circadian disruption on glucose metabolism: implications for type 2 diabetes. Diabetologia. 2020;63(3):462–472.

